# The phase of sensorimotor mu and beta oscillations has the opposite effect on corticospinal excitability

**DOI:** 10.1101/2022.02.22.481530

**Authors:** Miles Wischnewski, Zachary J. Haigh, Sina Shirinpour, Ivan Alekseichuk, Alexander Opitz

## Abstract

Neural oscillations in the primary motor cortex (M1) shape corticospinal excitability. Power and phase of ongoing mu (8-13 Hz) and beta (14-30 Hz) activity may mediate motor cortical output. However, the functional dynamics of both mu and beta phase and power relationships and their interaction, are largely unknown. Here, we employ recently developed real-time targeting of the mu and beta rhythm, to apply phase-specific brain stimulation and probe motor corticospinal excitability non-invasively. For this, we used instantaneous read-out and analysis of ongoing oscillations, targeting four different phases (0°, 90°, 180°, and 270°) of mu and beta rhythms with suprathreshold single-pulse transcranial magnetic stimulation (TMS) to M1. Ensuing motor evoked potentials (MEPs) in the right first dorsal interossei muscle were recorded. Twenty healthy adults took part in this double-blind randomized crossover study. Mixed model regression analyses showed significant phase-dependent modulation of corticospinal output by both mu and beta rhythm. Strikingly, these modulations exhibit a double dissociation. MEPs are larger at the mu trough and rising phase and smaller at the peak and falling phase. For the beta rhythm we found the opposite behavior. Also, mu power, but not beta power, was positively correlated with corticospinal output. Power and phase effects did not interact for either rhythm, suggesting independence between these aspects of oscillations. Our results provide insights into real-time motor cortical oscillation dynamics, which offers the opportunity to improve the effectiveness of TMS by specifically targeting different frequency bands.

## Introduction

Neocortical activity in the motor cortex is characterized by neural oscillations, foremost in the mu (8-13 Hz) and beta (14-30 Hz) rhythms. On the one hand, changes in their power correlate with motor functions such as preparation and execution of voluntary movement (Baker, 2007; Baker et al., 2003; Jenkinson & Brown, 2011; Jurkiewicz et al., 2006; Pfurtscheller et al., 1996; Pfurtscheller & Lopes Da Silva, 1999; Saleh et al., 2010). On the other hand, motor cortical output correlates with the phase of mu and beta oscillations (Berger et al., 2014; Combrisson et al., 2017; Miller et al., 2012; O’Keeffe et al., 2020; Yanagisawa et al., 2012). This phase-dependency may result from synchronization of neural spiking activity and is thus phase-specifically coupled to the oscillatory envelope (Fetz et al., 2000; Haegens et al., 2011; Johnson et al., 2020; Murthy & Fetz, 1992; 1996).

Although the coupling between cortical oscillation phase and spiking activity is well-established, how the phase of mu and beta oscillations in the motor cortex relates to functional cortical excitability is less clear. To provide causal evidence for a relation between oscillatory phase and cortical excitability, one needs to synchronize the electrocortical read-outs and causal probing of excitability with millisecond precision. Recent advances in real-time tracking of cortical oscillations and non-invasive modulation of motor cortex activity in healthy human participants have provided new insights (Bergmann et al., 2019; Madsen et al., 2019; Sasaki et al., 2021; Schaworonkow et al., 2018; 2019; Shirinpour et al., 2020; Zrenner et al., 2016; 2018). Such real-time systems, combining electroencephalography (EEG) and transcranial magnetic stimulation (TMS), have provided evidence for a modulation of corticospinal excitability by motor cortical oscillatory phase and power (Bergmann et al., 2019; Karabanov et al., 2021; Madsen et al., 2018; Zrenner et al., 2018).

Reports in non-human primates and patients with neurosurgical implants suggest that motor functioning is phase-dependent on oscillations in the motor cortical mu rhythm (Haegens et al., 2011; Yanagisawa et al., 2012). Based on this, first pursuits on real-time detection of motor oscillation phase relationships in healthy volunteers have focused on the mu rhythm (Bergmann et al., 2019; Madsen et al., 2019; Schaworonkow et al., 2018; 2019; Zrenner et al., 2018), Various studies suggest that motor evoked potential (MEP) amplitude is larger at the trough of the mu rhythm and smaller at the peak (Bergmann et al., 2019; Desideri et al., 2019; Schaworonkow et al., 2018; 2019; Zrenner et al., 2018). However, others have provided evidence that ongoing mu phase does not significantly predict corticospinal excitation (Karabanov et al., 2021; Madsen et al., 2019). Rather, pre-stimulation mu power is suggested to determine MEP amplitude (Bergmann et al., 2019; Karabanov et al., 2021; Madsen et al., 2019; Thies et al., 2020).

Whereas findings on associations between corticospinal excitability and mu phase are mixed, to the best of our knowledge, there are no real-time neuromodulation systems capable to target the beta rhythm non-invasively. Despite superficial similarities between mu and beta oscillations they reflect distinct functional sensorimotor networks and may have different anatomical origins (Gaetz & Cheyne, 2006; Jones et al., 2009; Premoli et al., 2017; Ronnqvist et al., 2013; Salmelin & Hari, 1994; Salmelin et al., 1995). As such, it is likely that phase-modulation of cortical excitability would reflect distinct patterns for mu and beta rhythms. Human and non-human primate studies have suggested a potential coupling of motor responses and motor cortical beta-phase (Miller et al., 2012; Murthy & Fetz, 1996; Reimer & Hatsopoulos, 2010). Electrocorticography (ECoG) has shown phase-dependency of motor network beta-rhythm activity in Parkinson’s disease patients (de Hemptine et al., 2013; Miller et al., 2012; O’Keeffe et al., 2020). Furthermore, beta phase-dependent stimulation in these patients has been shown to ameliorate motor deficits (Cagnan et al., 2017; Holt et al., 2019; Salimpour et al., 2022).

The absence of real-time TMS-EEG studies on beta rhythm may stem from the intrinsically lower signal-to-noise ratio, faster pace, and broader frequency band compared to mu oscillations. To reliably target the beta phase in real-time, we optimized a cutting-edge real-time algorithm -Educated Temporal Prediction (ETP) - to perform accurate forward predictions during real-time phase targeting (Shirinpour et al., 2020). Due to its robustness to noise and superior speed, ETP can accurately track and stimulate both mu and beta oscillations. Using our approach, we targeted mu and beta phase in the motor cortex in real-time. Our results show a double dissociation in the relationship between mu and beta phase on corticospinal excitability. That is, phases of mu oscillation that resulted in larger than average motor cortex output generate smaller than average motor cortex output for the same phases of beta, and vice versa. Our data provide the first evidence for distinct phase-dependency of mu-and beta-mediated functional sensorimotor networks that modulate corticospinal excitability. Optimizing TMS-targeting to mu or beta phase can increase robustness of TMS with clear implications for improving the efficacy of TMS in clinical use.

## Methods

### Participants

We recruited 20 healthy volunteers (11 female, mean ± std age: 22.7 y ± 2.9) in this double-blinded randomized crossover study. Each participant visited for two sessions (targeting mu and beta oscillations). Participants were right-handed, between 18 and 45 years of age, without a history of neurological or psychiatric disorders, head injuries, or metal or electric implants in the head, neck, or chest area. Participants were not pre-selected on the basis of electrophysiological characteristics, such as motor threshold or sensorimotor oscillatory power. The study was approved by the institutional review board of the University of Minnesota and all volunteers gave written informed consent prior to participation.

### Transcranial magnetic stimulation

We applied single-pulse biphasic TMS using the Magstim Rapid^2^ with a figure-of-eight shaped D70^2^ coil (Magstim Inc., Plymouth, MN, USA). The coil was placed over the left motor cortex, corresponding to the hotspot of the right first dorsal interossei (FDI) muscle, and oriented approximately at a 45° angle relative to the midline. Electromyography (EMG) was used to record motor-evoked potentials (MEP) from the FDI using self-adhesive, disposable electrodes. EMG sampling rate was set to 10 kHz using a BIOPAC ERS100C amplifier (BIOPAC systems, Inc., Goleta, CA, USA). Initially, the motor hotspot, i.e. the location and orientation that leads to the largest MEP, was determined. Hotspot coordinates were stored and coil location and orientation in reference to the hotspot were continuously tracked using a Brainsight neuronavigation system (Rogue Research Inc., Montreal, Canada). At the hotspot, the resting motor threshold (RMT) was determined using an adaptive threshold-hunting algorithm (Julkunen, 2019). The test intensity during the experimental session was set to 120% of RMT.

### EEG processing for real-time TMS triggering

Throughout the experiment, EEG was recorded using a 10-20 system, 64 active channel, TMS-compatible EEG system (actiCAP slim EEG cap, actiCHamp amplifier; Brain Products GmbH, Gilching, Germany). EEG data was streamed using Lab Streaming Layer (LSL) software to Matlab 2020b, where we used custom scripts to apply the ETP algorithm (Shirinpour et al., 2020). A sampling rate of 10 kHz with a 24-bits resolution per channel was used, and impedances were kept below 20 kΩ. The electrode of interest for this experiment was C3, located over the hand knob of the left sensorimotor area. To extract mu and beta oscillations unique to the electrode of interest, a Laplacian reference method was used, where the mean of the 8 surrounding electrodes was subtracted from the signal measured at C3 (Figure 1). This Laplacian C3 signal was used for real-time stimulation, as well as for offline analysis of mu and beta power.

**Figure 1.**
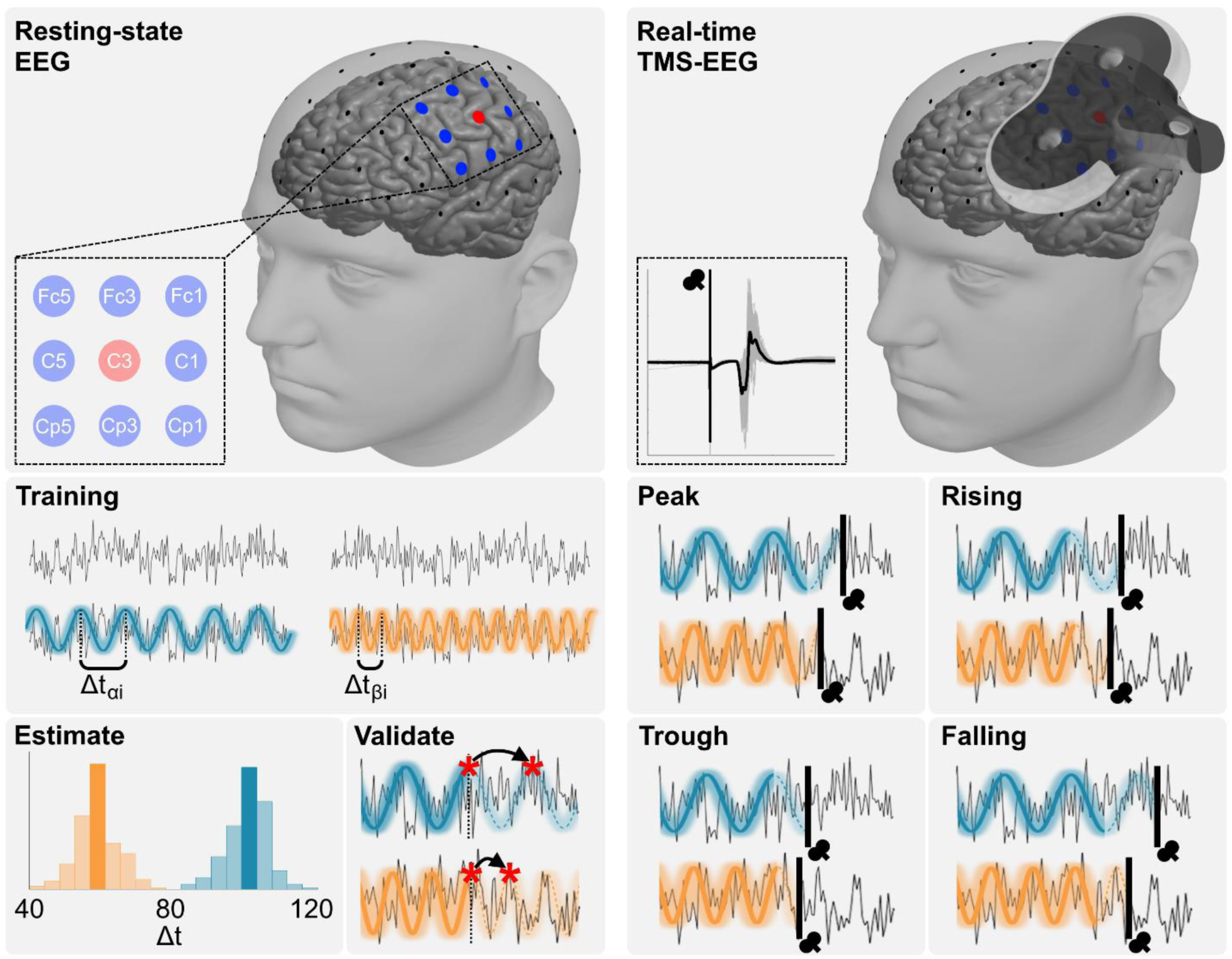
Overview of the educated temporal prediction (ETP) algorithm. Left: The algorithm is first trained using the resting state data from the sensorimotor cortex. Signals at sensorimotor cortex channel C3 are re-referenced using a center-surround Laplacian montage using 8 channels (Fc1, Fc3, Fc5, C1, C5, Cp1, Cp3, and Cp5). Depending on the experimental condition, we stimulated while tracking the phase of mu (8-13 Hz, blue) or beta (14-30 Hz, orange) range. From the resting-state data, the typical cycle length is extracted and used during the real-time stimulation. Right: During real-time application, EEG preprocessing follows the same pipeline as the training step. TMS is triggered at four different phases, namely peak (0°), rising phase (90°), trough (180°), or falling phase (270°). For each phase and oscillatory rhythm, we recorded MEPs from the FDI muscle.

The EEG-TMS setup for real-time stimulation used here follows our previously validated implementation (Shirinpour et al., 2020). In short, the ETP algorithm uses resting-state data from a training step before the real-time application, which provides an initial estimate of individual temporal dynamics of cortical oscillations. For this, we record resting-state data for three minutes perform a C3 Laplacian spatial filtering, and clean the signal using a zero-phase FIR (Finite Impulse Response) filter in the mu (8–13 Hz) or beta (14–30 Hz) range, as implemented in the Fieldtrip toolbox (Oostenveld et al 2010). Then, the algorithm estimates the typical cycle length (peak to peak interval) and validates its accuracy by simulating the accuracy of peak projection using the training data (Figure 1).

During real-time estimation, the calculated cycle length is adjusted to inform the forecasting algorithm that predicts upcoming peak, falling phase, trough, or rising phase (throughout this paper phase angles will be expressed in relation to a cosine, e.g. 0° is peak) of oscillation of interest and triggers TMS at the correct time. The EEG preprocessing pipeline during real-time measurements was the same as during the validation phase. Overall processing delay of our system, i.e. the time between sending trigger and actual pulse delivery was accounted for in our algorithm to accurately deliver the TMS at the desired phases (Shirinpour et al., 2020). Real-time TMS-EEG was performed in four blocks of 150 pulses. Within each block, phases were applied pseudorandomly. The experimenter and the participant were blinded to the phase order. A jittered interval between 2 and 3 seconds between consecutive triggers was introduced to minimize the direct effects of previous trials.

### Data processing and analysis

#### MEP analysis

We calculated peak-to-peak MEP amplitude using a custom Matlab script. MEPs were identified in a window between 20 and 60 ms after the TMS pulse. We excluded MEPs if average absolute EMG activity in a window from -100 to 0 ms before the TMS pulse was above 0.02 mV and larger than absolute average EMG activity + 2.5 times standard deviation in a window -500 to -400 ms before the TMS pulse (Wischnewski et al., 2016). All MEPs were visually inspected. Altogether, 3.3% of trials were removed (3.5% for targeting Mu phases and 3.0% for targeting Beta phases). For analysis, MEPs were normalized to the overall average.

#### Offline EEG analysis

Pre-TMS power was analyzed offline for inclusion in the main analysis. Raw EEG data were re-referenced to the Laplacian C3 montage as was used for online analyses (Figure 1). Data were epoched in a window between -1000 and 0 milliseconds with respect to TMS trigger and a bandpass filter (2-50 Hz) was applied. Pre-TMS power was calculated by applying a fast Fourier transform with Hanning taper at a resolution of 1 Hz as implemented in the Fieldtrip toolbox. Subsequently, we averaged power values between 8 and 13 Hz (mu power) and between 14 and 30 Hz (beta power) at the single-trial level.

To investigate potential differences in mu and beta oscillation topography, sensor-level distributions were examined. Resting-state EEG data were re-referenced to Cz and filtered in the mu (8-13 Hz) and beta (14-30 Hz) bands, respectively. We estimated the pairwise correlations between the electrode of interest C3 to all other electrodes. Topographic plots were used to depict the spatial distribution of the correlations for mu and beta separately, as well as the difference between both conditions.

#### Statistical analysis

In a trial level analysis, a general linear mixed-effects model (GLMM) was used on trial data with target phase (peak, falling, trough, rising) and target rhythm (mu, beta) as fixed effects variable and participant number as random effects variable. MEP amplitude was the dependent variable. Independently, after averaging MEPs per phase for each participant, Raleigh’s Z test of non-uniformity was performed for phase modulation at Mu and Beta oscillations.

To test the effects of pre-TMS power, GLMMs were run on Mu and Beta conditions separately adding respective pre-TMS power as a continuous fixed effects variable. These analyses were followed up by post hoc subject-level simple linear regression models. Subsequently, Spearman rank correlation between pre-TMS power and MEP amplitude for each subject and session were calculated.

Finally, Spearman rank correlation was performed on the topographic distribution of mu and beta oscillations. For all analyses, significance level was set at α = 0.05.

## Results

Real-time TMS of ongoing cortical oscillations resulted in a double dissociation of phase relationships for Mu and Beta oscillations (Figure 2A). Accordingly, GLMM regression showed a significant interaction between target phase and target rhythm on MEP amplitude (F = 16.42, p < 0.001). Distinct phase relation patterns were confirmed by Rayleigh’s test for non-uniformity of circular group level data. Normalized MEP amplitudes at phases of the Mu rhythm were non-uniformly distributed (Z = 3.02, p = 0.048), with a mean direction of the circular distribution of *θ* = 225.00° and circular dispersion of *κ* = 29.27°. Thus, MEP amplitudes were maximal when Mu oscillations are at trough and rising phase (Figure 2B) and lower than average at the opposing phases. Normalized MEP amplitudes at phases of the Beta rhythm were also non-uniformly distributed (Z = 3.27, p = 0.037), with circular mean of *θ* = 29.05° and dispersion of *κ* = 30.53°. This means that MEP amplitudes were maximal when beta oscillations are at peak or falling phase (Figure 2B) and again lower than average at the opposing phases.

**Figure 2.**
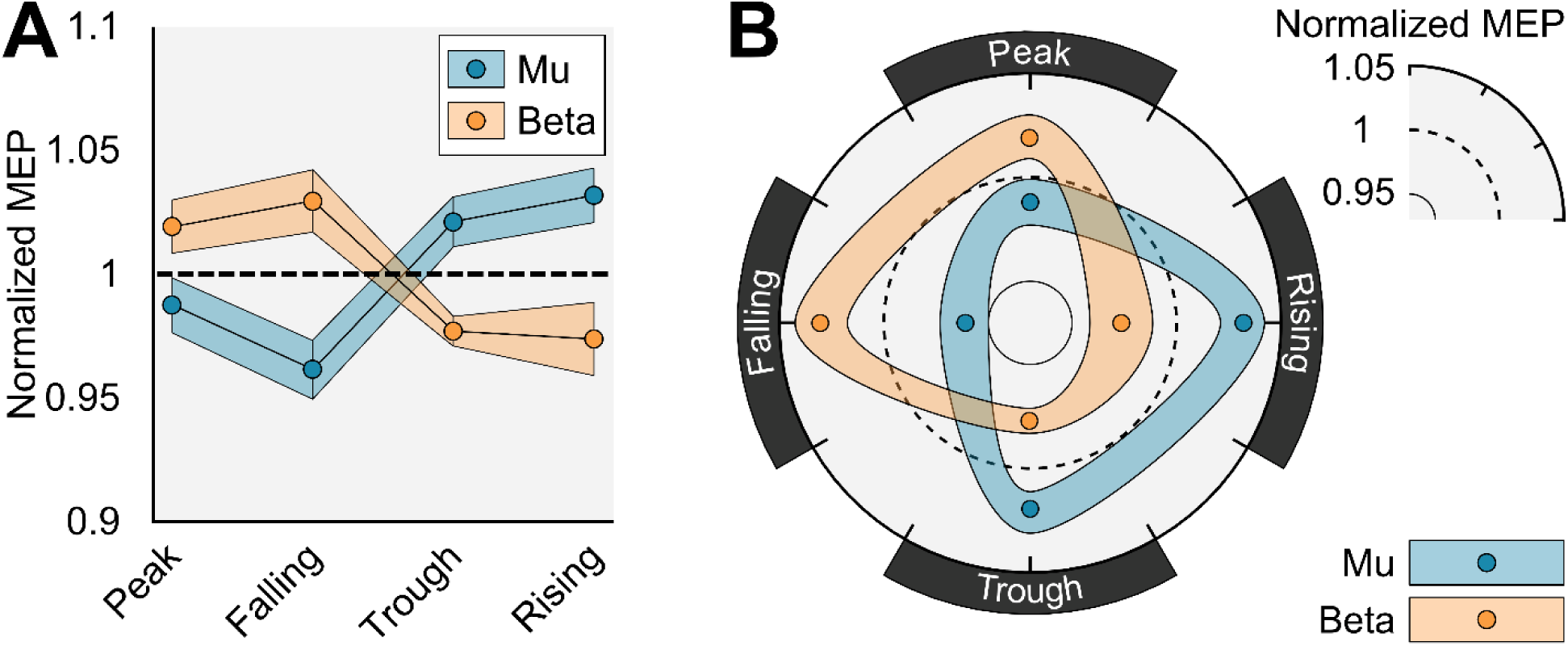
A) Group average (n = 20) ± standard error of mean of normalized MEPs for targeted phases in the mu and beta frequency. B) Circular representation of the data with smooth interpolation between conditions.

The results are largely consistent at the individual level. The observed pattern of larger MEP amplitudes at the beta peak compared to the mu peak were observed in 13 out of 20 participants. Larger MEP amplitudes at beta falling compared to mu falling were observed in 14 out of 20 participants. Larger MEP amplitudes at mu trough compared to beta trough were observed in 18 out of 20 participants. Larger MEP amplitudes at mu rising compared to beta rising were observed in 14 out of 20 participants (Figure 3).

**Figure 3.**
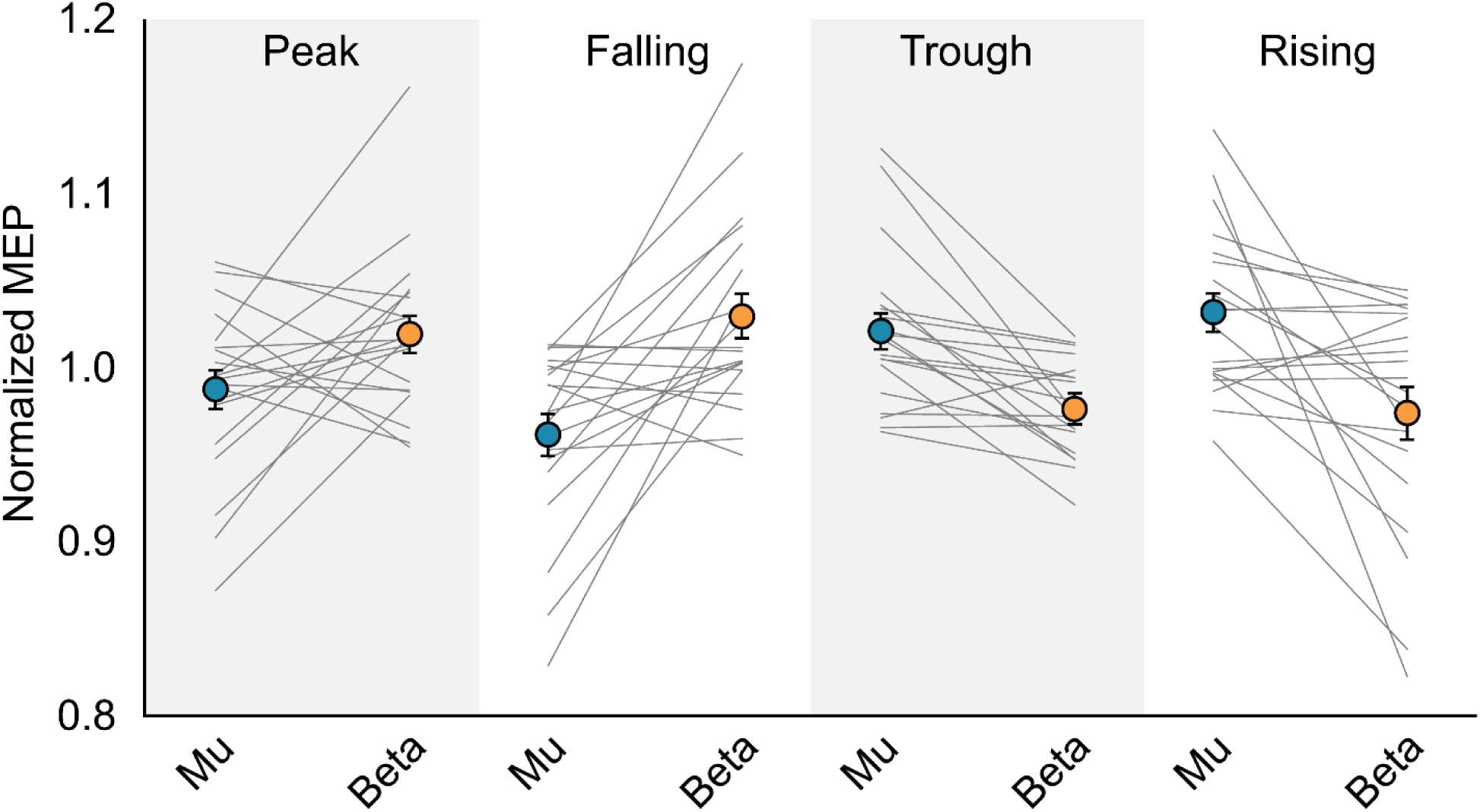
Individual phase-dependent modulation of MEP amplitude for mu and beta oscillations. Error bars represent standard error of mean.

In analyses of each target rhythm condition separately, we added pre-TMS power of the targeted rhythm. MEP amplitude during targeting of the Mu rhythm was affected by both target phase (F = 3.52, p = 0.014) and pre-TMS mu power (F = 11.99, p = 0.001). Crucially, however, no significant phase*power interaction was observed (F = 1.65, p = 0.175), suggesting that both power and phase affect MEP amplitude independently. At an individual level, correlation between Mu power and MEP amplitude ranged between ρ = -0.111 and ρ = 0.343 (median ρ = 0.071). A significant positive relationship was observed in 15 out of 40 sessions, whereas a significant negative relationship was observed in 1 session (Figure 4A). MEP amplitude while targeting beta rhythm was affected by target phase alone (F = 2.79, p = 0.038). No effect of pre-TMS beta power (F = 2.06, p = 0.151), nor a phase*power interaction (F = 2.16, p = 0.091) was observed on MEP amplitude. At an individual level, correlation between Beta power and MEP amplitude ranged between ρ = -0.161 and ρ = 0.267 (median ρ = 0.053). A significant positive relationship was observed in 12 out of 40 sessions, whereas a significant negative relationship was observed in 4 out of 40 sessions (Figure 4B).

**Figure 4.**
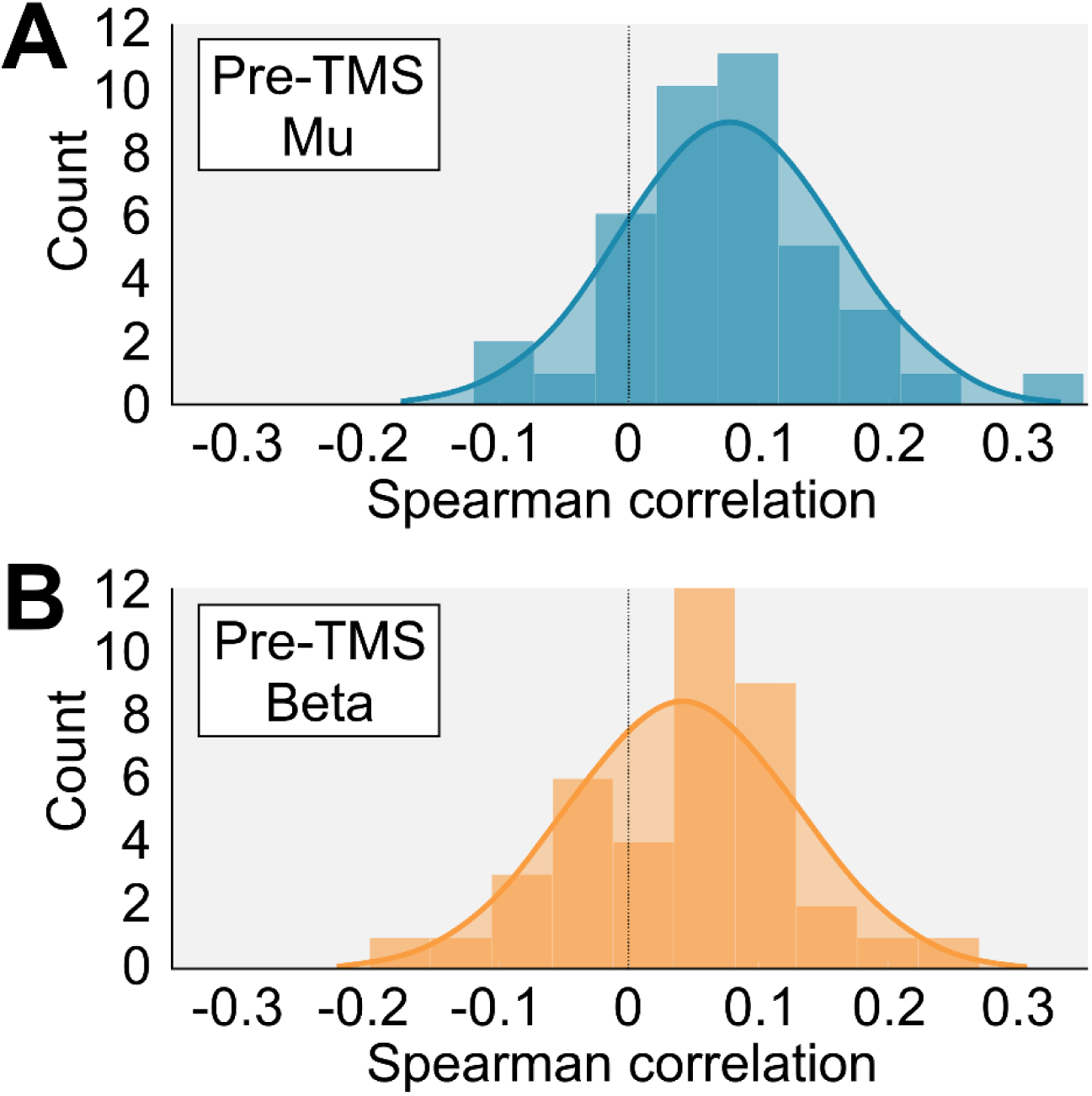
Histogram of individual Spearman correlations between MEP amplitude and A) pre-TMS mu power, and B) pre-TMS beta power.

One possible confound could arise where channels in the Laplacian reference montage contribute differently to the target electrode between conditions. Therefore, we performed a sensor-level analysis of mu and beta distributions, by looking at the channel-to-channel correlations. Resulting topographic plots showed highly similar distributions for both mu and beta rhythms at sensor level (Figure 5). Distributions were highly correlated (ρ = 0.975, p < 0.001), suggesting that our main results cannot be explained by differences in mu and beta signal arrangement.

**Figure 5.**
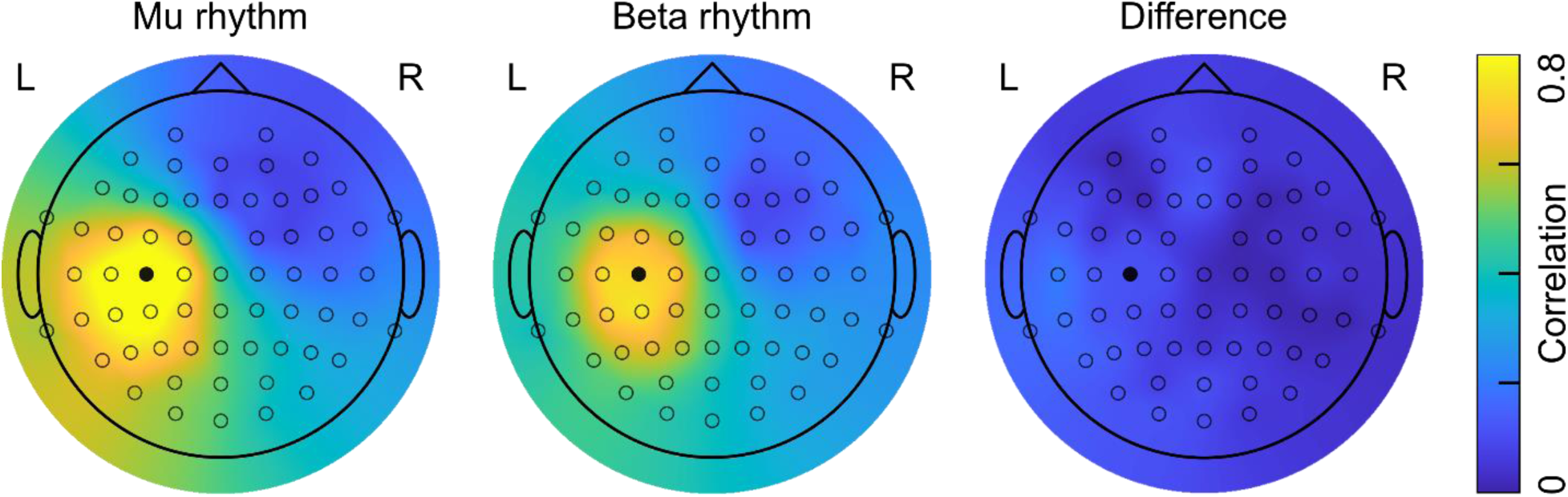
Spatial topographies for the recorded mu rhythm, beta rhythm, and the difference between both. Color map represents correlational values of electrode pairings between target electrode C3 and all other electrodes. The black electrode corresponds to C3.

## Discussion

In this study, we demonstrate for the first time that mu and beta oscillation phase differentially modulate MEP amplitude. In summary, we found that I) phase of mu and beta oscillations picked up at sensorimotor channels modulate corticospinal excitation; II) this phase-dependent MEP modulation follows an opposing pattern for mu and beta; III) mu power, but not beta power, significantly modulates MEP amplitude; IV) modulation of MEP amplitudes by phase and power do not interact.

To our knowledge, we provide the first direct evidence for MEP amplitude modulation by beta phase, in addition to mu phase, measured with real-time TMS-EEG. Beta-phase dependency has been hinted at by previous offline TMS studies using post-hoc analyses (Hussain et al., 2019; Keil et al., 2014; Khademi et al., 2017; Schilberg et al., 2021; Torrecillos et al., 2020; van Elswijk et al., 2010). Also, human subdural electrocorticographic (ECoG) recordings have shown that motor cortical beta activity is phase-locked to neural population activity during movement (de Hemptinne et al., 2013; Holt et al., 2019; Miller et al., 2012). Furthermore, motor cortical spiking activity has been shown to be dependent on local field potential beta-phase in non-human primates (Reimer & Hatsopoulos, 2010; Witham et al., 2007). Sensorimotor beta oscillations have been suggested to arise from alternating de-and hyper-polarization of layer V pyramidal cells, mediated by phase-locked gamma-aminobutyric acid (GABA) mediated interneuron inputs (Baker, 2007; Bhatt et al., 2016; Lacey et al., 2014; Rossiter et al., 2014).

Here we show that beta phase-dependency can be probed non-invasively in real-time. Our data showed largest MEP amplitudes during beta peak and falling phase (Figure 2). Salimpour et al. (2022) applied real-time electrical motor cortex stimulation in Parkinson’s disease patients during surgery. Although direct comparison of results from electrical stimulation and ECoG data to ours may be challenging, it is interesting to point out that phase-dependency was similar, with beta peak and falling phase leading to the largest motor output.

We found no dependency of beta power on MEP amplitude, nor was there an interaction between beta phase and power, in line with previous findings (Hussain et al., 2019; Mitchell et al., 2007; Ogata et al., 2019; Peters et al., 2020; Schilberg et al., 2021). This should not imply that beta oscillations are not related to motor output and evidence from previous research suggests that the relationship between beta oscillations and motor activation is complex. Pre-movement reduction of beta power has been associated with faster voluntary movement (Khanna & Carmena, 2017). Chronic elevation of beta power, observed in Parkinson’s disease has been related to difficulty initiating and controlling movements (Brown, 2006; Cannon et al., 2014; Kühn et al., 2006). Furthermore, in addition to low-amplitude ongoing beta activity, high-amplitude beta bursts are suggested to be positively correlated to movement control (Bonaiuto et al., 2021; Chen & Fetz, 2005; Feingold et al., 2015; O’Keeffe et al., 2020; Reimer & Hatsopoulos, 2010). Although these behavioral studies imply that beta power and beta bursts are crucial for endogenous control of voluntary movement, our and previous studies suggest that they are not related to exogenously probed cortico-spinal excitability (Hussain et al., 2019; Ogata et al., 2019, Peters et al., 2020). Furthermore, Peters and colleagues (2020) found that pre-TMS resting beta power does not affect the propagation of TMS excitations throughout the cortical-subcortical motor network. Therefore, it seems that beta power may be a predictor for corticospinal activation during voluntary or task-related motor control, but not during resting-state motor excitability per se.

Additionally, we found that corticospinal excitation was modulated by the mu rhythm with an opposite phase relationship compared to beta oscillations. Various studies previously indicated mu phase-dependent modulation of MEP amplitudes, with larger responses at the mu trough compared to mu peak. (Bergmann et al., 2019; Desideri et al., 2019; Schaworonkow et al., 2018; 2019; Zrenner et al., 2018). By real-time targeting of four, rather than two phases, our results extend previous findings by showing that in particular the trough and subsequent rising phase yield largest corticospinal excitation, whereas mu peak and falling yield the smallest motor cortex activation (Figure 2).

Pre-stimulus mu power was a significant predictor for corticospinal excitability, but did not interact with mu phase, suggesting independence between mu power and phase. Subject-level positive correlations were observed in majority of subjects. Although the observed relationship was relatively weak -correlations varying between -0.1 and 0.3 - it is in line with previous observations (Bergmann et al., 2019; Karabanov et al., 2021; Schilberg et al., 2021; Thies et al., 2020). However, others have found no relationship between mu power and MEP amplitude (Berger et al., 2014; Zrenner et al., 2018), or even a negative association (Madsen et al., 2019; Sauseng et al., 2009, Zarkowski et al., 2006). At a first glance, a positive relationship between mu power and corticospinal activity seems counterintuitive since sensorimotor mu oscillations are related to GABAa-mediated inhibitory activity (Bergmann et al., 2019). Also, higher mu power has been shown to reduce TMS-induced blood oxygenation level-dependent (BOLD) responses throughout the cortical-subcortical motor network (Peters et al., 2020). However, mu oscillations are thought to predominantly originate from the somatosensory cortex (Gaetz & Cheyne, 2006; Jones et al., 2009; Premoli et al., 2017; Ronnqvist et al., 2013; Salmelin & Hari, 1994; Salmelin et al., 1995). Interconnections between somatosensory and primary motor cortex comprise of an intricate network of excitatory and inhibitory reciprocal connections. Increased mu power may reflect feedforward inhibition to primary motor cortex resulting in local disinhibition (Thies et al., 2020). This may result in a net-facilitation or net-inhibition of corticospinal activation, yielding contradictory findings between studies.

Sensorimotor mu and beta oscillations have been suggested to stem from distinct neural origins (Gaetz & Cheyne, 2006; Jones et al., 2009; Premoli et al., 2017; Ronnqvist et al., 2013; Salmelin & Hari, 1994; Salmelin et al., 1995). Specifically, mu oscillations are proposed to originate pre-dominantly from the post-central gyrus (Salmelin & Hari, 1994; Salmelin et al., 1995), although pre-central origins of mu have been reported as well (Ronnqvist et al., 2013; Smit et al., 2013; Szurhaj et al., 2003). In contrast, beta oscillations are thought to stem from pre-central primary motor cortex (Donoghue et al., 1998; Jurkiewicz et al., 2006; Salmelin & Hari, 1994; Salmelin et al., 1995), but are also observed in post-central somatosensory cortex (Brovelli et al., 2004; Jurkiewicz et al., 2006; Szurhaj et al., 2003). Although our study cannot make inferences on the source of mu and beta oscillations, sensor-level signal distributions were highly similar (Figure 5). Similar scalp-level topographies suggest that potential differences in neural origin did not influence phase detection during real-time stimulation. A potential explanation for the opposing phase-relationship we observed results from differences in axonal orientation within mu and beta sources. This possibility could be investigated in future studies.

Our findings are crucial for the improvement of TMS effectiveness for treatment of neurological and psychiatric disorders. Targeting optimal rhythms with repetitive TMS could decrease variability of TMS outcomes (Baur et al., 2020; Zrenner et al., 2018). For instance, targeting optimal oscillation phase could improve efficacy of TMS in the recovery of stroke (Hussain et al., 2020) and treatment of major depressive disorder (Zrenner et al., 2020). In this study, to our knowledge, we were able to non-invasively target the beta rhythm in real-time reliably for the first time. In future work it will be crucial to further optimize real-time and closed-loop systems, in order to target different oscillatory rhythms, and different spatial locations (Metsomaa et al., 2021). Eventually, this will allow for adaptive non-invasive neuromodulation that can provide personalized decoding of on-going brain states. This individualization can greatly benefit clinical application of TMS, by reducing variability between and within patients.

## Notes

### Competing Interest Statement

The authors have declared no competing interest.

## References

Baker S.N. (2007). Oscillatory interactions between sensorimotor cortex and the periphery. Curr. Opin. Neurobiol. 17(6), 649–655. https://doi.org/10.1016/j.conb.2008.01.007

Baker, S.N., Pinches, E.M., Lemon, R.N. (2003). Synchronization in monkey motor cortex during a precision grip task. II. effect of oscillatory activity on corticospinal output. J. Neurophysiol. 89(4), 1941–1953. https://doi.org/10.1152/jn.00832.2002

Baur, D., Galevska, D., Hussain, S., Cohen, L. G., Ziemann, U., Zrenner, C. (2020). Induction of LTD-like corticospinal plasticity by low-frequency rTMS depends on pre-stimulus phase of sensorimotor μ-rhythm. Brain Stimul. 13(6), 1580–1587. https://doi.org/10.1016/j.brs.2020.09.005

Berger, B., Minarik, T., Liuzzi, G., Hummel, F.C., Sauseng, P. (2014). EEG oscillatory phase-dependent markers of corticospinal excitability in the resting brain. Biomed. Res. Int. 2014, 936096. https://doi.org/10.1155/2014/936096

Bergmann, T.O., Lieb, A., Zrenner, C., Ziemann, U. (2019). Pulsed Facilitation of Corticospinal Excitability by the Sensorimotor μ-Alpha Rhythm. J. Neurosci. 39(50), 10034–10043. https://doi.org/10.1523/JNEUROSCI.1730-19.2019

Bhatt, M.B., Bowen, S., Rossiter, H.E., Dupont-Hadwen, J., Moran, R.J., Friston, K.J., Ward, N.S. (2016). Computational modelling of movement-related beta-oscillatory dynamics in human motor cortex. NeuroImage 133, 224–232. https://doi.org/10.1016/j.neuroimage.2016.02.078

Bonaiuto, J. J., Little, S., Neymotin, S. A., Jones, S. R., Barnes, G. R., & Bestmann, S. (2021). Laminar dynamics of high amplitude beta bursts in human motor cortex. NeuroImage, 242, 118479. https://doi.org/10.1016/j.neuroimage.2021.118479

Brovelli, A., Ding, M., Ledberg, A., Chen, Y., Nakamura, R., Bressler, S.L. (2004). Beta oscillations in a large-scale sensorimotor cortical network: directional influences revealed by Granger causality. Proc. Nat. Acad. Sci. U. S. A. 101(26), 9849–9854. https://doi.org/10.1073/pnas.0308538101

Brown P. (2006). Bad oscillations in Parkinson’s disease. J. Neural Transm. (70), 27–30. https://doi.org/10.1007/978-3-211-45295-0_6

Cagnan, H., Pedrosa, D., Little, S., Pogosyan, A., Cheeran, B., Aziz, T. (2017). Stimulating at the right time: phase-specific deep brain stimulation. Brain 140(1), 132–145. https://doi.org/10.1093/brain/aww286

Cannon, J., McCarthy, M. M., Lee, S., Lee, J., Börgers, C., Whittington, M. A., Kopell, N. (2014). Neurosystems: brain rhythms and cognitive processing. Eur. J. Neurosci. 39(5), 705–719. https://doi.org/10.1111/ejn.12453

Chen, D., Fetz, E.E. (2005). Characteristic membrane potential trajectories in primate sensorimotor cortex neurons recorded in vivo. J. Neurophysiol. 94(4), 2713–2725. https://doi.org/10.1152/jn.00024.2005

Combrisson, E., Perrone-Bertolotti, M., Soto, J.L., Alamian, G., Kahane, P., Lachaux, J.P., Guillot, A., Jerbi, K. (2017). From intentions to actions: Neural oscillations encode motor processes through phase, amplitude and phase-amplitude coupling. NeuroImage 147, 473–487. https://doi.org/10.1016/j.neuroimage.2016.11.042

de Hemptinne, C., Ryapolova-Webb, E.S., Air, E.L., Garcia, P.A., Miller, K.J., Ojemann, J.G., et al. (2013). Exaggerated phase-amplitude coupling in the primary motor cortex in Parkinson disease. Proc. Nat. Acad. Sci. U. S. A. 110(12), 4780–4785. https://doi.org/10.1073/pnas.1214546110

Desideri, D., Zrenner, C., Ziemann, U., Belardinelli, P. (2019). Phase of sensorimotor μ-oscillation modulates cortical responses to transcranial magnetic stimulation of the human motor cortex. J. Physiol. 597(23), 5671–5686. https://doi.org/10.1113/JP278638

Donoghue, J.P., Sanes, J.N., Hatsopoulos, N.G., Gaál, G. (1998). Neural discharge and local field potential oscillations in primate motor cortex during voluntary movements. J. Neurophysiol. 79(1), 159–173. https://doi.org/10.1152/jn.1998.79.1.159

Fetz, E.E., Chen, D., Murthy, V.N., Matsumura, M. (2000). Synaptic interactions mediating synchrony and oscillations in primate sensorimotor cortex. J. Phsyiol Paris, 94(5-6), 323–331. https://doi.org/10.1016/s0928-4257(00)01089-5

Feingold, J., Gibson, D.J., DePasquale, B., Graybiel, A.M. (2015). Bursts of beta oscillation differentiate postperformance activity in the striatum and motor cortex of monkeys performing movement tasks. Proc. Nat. Acad. Sci. U.S.A. 112(44), 13687–13692. https://doi.org/10.1073/pnas.1517629112

Gaetz, W., Cheyne, D. (2006). Localization of sensorimotor cortical rhythms induced by tactile stimulation using spatially filtered MEG. NeuroImage, 30(3), 899–908. https://doi.org/10.1016/j.neuroimage.2005.10.009

Haegens, S., Nacher, V., Luna, R., Romo, R., Jensen, O. (2011). α-Oscillations in the monkey sensorimotor network influence discrimination performance by rhythmical inhibition of neuronal spiking. Proc. Natl. Acad. Sci. 108(48), 19377–19382. https://doi.org/10.1073/pnas.1117190108

Holt, A.B., Kormann, E., Gulberti, A., Pötter-Nerger, M., McNamara, C.G., Cagnan, H., et al. (2019). Phase-Dependent Suppression of Beta Oscillations in Parkinson’s Disease Patients. J. Neurosci. 39(6), 1119–1134. https://doi.org/10.1523/JNEUROSCI.1913-18.2018

Hussain, S. J., Claudino, L., Bönstrup, M., Norato, G., Cruciani, G., Thompson, R., Zrenner, C., Ziemann, U., Buch, E., Cohen, L. G. (2019). Sensorimotor Oscillatory Phase-Power Interaction Gates Resting Human Corticospinal Output. Cereb Cortex 29(9), 3766–3777. https://doi.org/10.1093/cercor/bhy255

Hussain, S.J., Hayward, W., Fourcand, F., Zrenner, C., Ziemann, U., Buch, E.R., et al. (2020) Phase-dependent transcranial magnetic stimulation of the lesioned hemisphere is accurate after stroke. Brain Stimul. 13(5), 1354–1357. https://doi.org/10.1016/j.brs.2020.07.005

Jenkinson, N., Brown, P. (2011). New insights into the relationship between dopamine, beta oscillations and motor function. Trends Neurosci. 34(12), 611–618. https://doi.org/10.1016/j.tins.2011.09.003

Johnson, L., Alekseichuk, I., Krieg, J., Doyle, A., Yu, Y., Vitek, J., Johnson, M., Opitz, A. (2020). Dose-dependent effects of transcranial alternating current stimulation on spike timing in awake nonhuman primates. Sci. Adv. 6(36), eaaz2747. https://doi.org/10.1126/sciadv.aaz2747

Jones, S.R., Pritchett, D.L., Sikora, M.A., Stufflebeam, S.M., Hämäläinen, M., Moore, C.I. (2009). Quantitative analysis and biophysically realistic neural modeling of the MEG mu rhythm: rhythmogenesis and modulation of sensory-evoked responses. J. Neurophysiol. 102(6), 3554–3572. https://doi.org/10.1152/jn.00535.2009

Julkunen P. (2019). Mobile Application for Adaptive Threshold Hunting in Transcranial Magnetic Stimulation. IEEE Trans. Neural. Syst. Rehabil. Eng. 27(8), 1504–1510. https://doi.org/10.1109/TNSRE.2019.2925904

Jurkiewicz, M.T., Gaetz, W.C., Bostan, A.C., Cheyne, D. (2006). Post-movement beta rebound is generated in motor cortex: evidence from neuromagnetic recordings. NeuroImage, 32(3), 1281–1289. https://doi.org/10.1016/j.neuroimage.2006.06.005

Karabanov, A.N., Madsen, K.H., Krohne, L. G., Siebner, H.R. (2021). Does pericentral mu-rhythm “power” corticomotor excitability? - A matter of EEG perspective. Brain stimulation, 14(3), 713–722. https://doi.org/10.1016/j.brs.2021.03.017

Keil, J., Timm, J., Sanmiguel, I., Schulz, H., Obleser, J., Schönwiesner, M. (2014). Cortical brain states and corticospinal synchronization influence TMS-evoked motor potentials. J. Neurophysiol. 111(3), 513–519. https://doi.org/10.1152/jn.00387.2013

Khademi, F., Royter, V., Gharabaghi, A. (2018). Distinct Beta-band Oscillatory Circuits Underlie Corticospinal Gain Modulation. Cereb. Cortex 28(4), 1502–1515. https://doi.org/10.1093/cercor/bhy016

Khanna, P., & Carmena, J. M. (2017). Beta band oscillations in motor cortex reflect neural population signals that delay movement onset. eLife, 6, e24573. https://doi.org/10.7554/eLife.24573

Kühn, A.A., Kupsch, A., Schneider, G.H. Brown, P. (2006). Reduction in subthalamic 8–25 Hz oscillatory activity correlates with clinical improvement in Parkinson’s disease. Eur. J. Neurosci. 23, 1956– 1960. https://doi.org/10.1111/j.1460-9568.2006.04717.x

Madsen, K.H., Karabanov, A.N., Krohne, L.G., Safeldt, M.G., Tomasovic, L., Siebner, H.R. (2019). No trace of phase: Corticomotor excitability is not tuned by phase of pericentral mu-rhythm. Brain Stimul. 12(5), 1261–1270. https://doi.org/10.1016/j.brs.2019.05.005

Metsomaa, J., Belardinelli, P., Ermolova, M., Ziemann, U., Zrenner, C. (2021). Causal decoding of individual cortical excitability states. NeuroImage, 245, 118652. https://doi.org/10.1016/j.neuroimage.2021.118652

Miller, K.J., Hermes, D., Honey, C.J., Hebb, A.O., Ramsey, N.F., Knight, R.T., Ojemann, J.G., Fetz, E.E. (2012). Human motor cortical activity is selectively phase-entrained on underlying rhythms. PLoS Comput. Biol, 8(9), e1002655. https://doi.org/10.1371/journal.pcbi.1002655

Mitchell, W.K., Baker, M.R., Baker, S.N. (2007). Muscle responses to transcranial stimulation in man depend on background oscillatory activity. J. Physiol. 583(Pt 2), 567–579. https://doi.org/10.1113/jphysiol.2007.134031

Murthy, V.N., Fetz, E.E. (1992). Coherent 25-to 35-Hz oscillations in the sensorimotor cortex of awake behaving monkeys. Proc. Nat. Acad. Sci. U. S. A. 89(12), 5670–5674. https://doi.org/10.1073/pnas.89.12.5670

Murthy, V. N., Fetz, E.E. (1996). Synchronization of neurons during local field potential oscillations in sensorimotor cortex of awake monkeys. J. Neurophysiol. 76(6), 3968– 3982. https://doi.org/10.1152/jn.1996.76.6.3968

Ogata, K., Nakazono, H., Uehara, T., Tobimatsu, S. (2019). Prestimulus cortical EEG oscillations can predict the excitability of the primary motor cortex. Brain Stimul. 12(6), 1508–1516. https://doi.org/10.1016/j.brs.2019.06.013

O’Keeffe, A. B., Malekmohammadi, M., Sparks, H., Pouratian, N. (2020). Synchrony drives motor cortex beta bursting, waveform dynamics, and phase-amplitude coupling in Parkinson’s disease. J. Neurosci. 40(30), 5833–5846. https://doi.org/10.1523/JNEUROSCI.1996-19.2020

Peters, J.C., Reithler, J., Graaf, T.A., Schuhmann, T., Goebel, R., Sack, A.T. (2020). Concurrent human TMS-EEG-fMRI enables monitoring of oscillatory brain state-dependent gating of cortico-subcortical network activity. Commun. Biol. 3(1), 40. https://doi.org/10.1038/s42003-020-0764-0

Pfurtscheller, G., Lopes da Silva, F.H. (1999). Event-related EEG/MEG synchronization and desynchronization: basic principles. Clin. Neurophysiol. 110(11), 1842–1857. https://doi.org/10.1016/s1388-2457(99)00141-8

Pfurtscheller, G., Stancák, A., Jr, Neuper, C. (1996). Post-movement beta synchronization. A correlate of an idling motor area?. Electroencephal. Clin. Neurophysiol. 98(4), 281–293. https://doi.org/10.1016/0013-4694(95)00258-8

Premoli, I., Bergmann, T.O., Fecchio, M., Rosanova, M., Biondi, A., Belardinelli, P., Ziemann, U. (2017). The impact of GABAergic drugs on TMS-induced brain oscillations in human motor cortex. NeuroImage 163, 1–12. https://doi.org/10.1016/j.neuroimage.2017.09.023

Reimer, J., Hatsopoulos, N.G. (2010). Periodicity and evoked responses in motor cortex. J. Neurosci. 30(34), 11506–11515. https://doi.org/10.1523/JNEUROSCI.5947-09.2010

Ronnqvist, K.C., McAllister, C.J., Woodhall, G.L., Stanford, I.M., Hall, S.D. (2013). A multimodal perspective on the composition of cortical oscillations. Front. Hum. Neurosci. 7, 132. https://doi.org/10.3389/fnhum.2013.00132

Rossiter, H.E., Davis, E.M., Clark, E.V., Boudrias, M.H., Ward, N.S. (2014). Beta oscillations reflect changes in motor cortex inhibition in healthy ageing. NeuroImage 91(100), 360– 365. https://doi.org/10.1016/j.neuroimage.2014.01.012

Saleh, M., Reimer, J., Penn, R., Ojakangas, C.L., Hatsopoulos, N.G. (2010). Fast and slow oscillations in human primary motor cortex predict oncoming behaviorally relevant cues. Neuron 65(4), 461–471. https://doi.org/10.1016/j.neuron.2010.02.001

Salimpour, Y., Mills, K.A., Hwang, B.Y., Anderson, W.S. (2022). Phase-targeted stimulation modulates phase-amplitude coupling in the motor cortex of the human brain. Brain Stimul. 15(1), 152–163. https://doi.org/10.1016/j.brs.2021.11.019

Salmelin, R., Hari, R. (1994). Spatiotemporal characteristics of sensorimotor neuromagnetic rhythms related to thumb movement. Neurosci. 60(2), 537–550. https://doi.org/10.1016/0306-4522(94)90263-1

Salmelin, R., Hämäläinen, M., Kajola, M., Hari, R. (1995). Functional segregation of movement-related rhythmic activity in the human brain. NeuroImage 2(4), 237–243. https://doi.org/10.1006/nimg.1995.1031

Sasaki, K., Fujishige, Y., Kikuchi, Y., Odagaki, M. (2021). A transcranial magnetic stimulation trigger system for suppressing motor-evoked potential fluctuation using electroencephalogram coherence analysis: algorithm development and validation study JMIR Biomed. Eng. 6(2), e28902. https://doi.org/10.2196/28902

Sauseng, P., Klimesch, W., Gerloff, C., Hummel, F.C. (2009). Spontaneous locally restricted EEG alpha activity determines cortical excitability in the motor cortex. Neuropsychologia 47(1), 284–288. https://doi.org/10.1016/j.neuropsychologia.2008.07.021

Schaworonkow, N., Caldana Gordon, P., Belardinelli, P., Ziemann, U., Bergmann, T.O., Zrenner, C. (2018). μ-Rhythm Extracted With Personalized EEG Filters Correlates With Corticospinal Excitability in Real-Time Phase-Triggered EEG-TMS. Front. Neurosci. 12, 954. https://doi.org/10.3389/fnins.2018.00954

Schaworonkow, N., Triesch, J., Ziemann, U., Zrenner, C. (2019). EEG-triggered TMS reveals stronger brain state-dependent modulation of motor evoked potentials at weaker stimulation intensities. Brain Stimul. 12(1), 110–118. https://doi.org/10.1016/j.brs.2018.09.009

Schilberg, L., Ten Oever, S., Schuhmann, T., Sack, A.T. (2021). Phase and power modulations on the amplitude of TMS-induced motor evoked potentials. PloS one 16(9), e0255815. https://doi.org/10.1371/journal.pone.0255815

Shirinpour, S., Alekseichuk, I., Mantell, K., Opitz, A. (2020). Experimental evaluation of methods for real-time EEG phase-specific transcranial magnetic stimulation. J. Neural Eng. 17(4), 046002. https://doi.org/10.1088/1741-2552/ab9dba

Smit, D.J., Linkenkaer-Hansen, K., de Geus, E.J. (2013). Long-range temporal correlations in resting-state α oscillations predict human timing-error dynamics. J. Neurosci. (27), 11212–11220. https://doi.org/10.1523/JNEUROSCI.2816-12.2013

Stefanou, M.I., Baur, D., Belardinelli, P., Bergmann, T.O., Blum, C., Gordon, P.C., et al. (2019). Brain state-dependent brain stimulation with real-time electroencephalography-triggered transcranial magnetic stimulation. Journal of visualized experiments. JoVE (150), 10.3791/59711. https://doi.org/10.3791/59711

Szurhaj, W., Derambure, P., Labyt, E., Cassim, F., Bourriez, J. L., Isnard, J., et al. (2003). Basic mechanisms of central rhythms reactivity to preparation and execution of a voluntary movement: a stereoelectroencephalographic study. Clin. Neurophysiol. 114(1), 107–119. https://doi.org/10.1016/s1388-2457(02)00333-4

Thies, M., Zrenner, C., Ziemann, U., Bergmann, T.O. (2018). Sensorimotor mu-alpha power is positively related to corticospinal excitability. Brain Stimul. 11(5), 1119–1122. https://doi.org/10.1016/j.brs.2018.06.006

Torrecillos, F., Falato, E., Pogosyan, A., West, T., Di Lazzaro, V., Brown, P. (2020). Motor cortex inputs at the optimum phase of beta cortical oscillations undergo more rapid and less variable corticospinal propagation. J. Neursci. 40(2), 369–381. https://doi.org/10.1523/JNEUROSCI.1953-19.2019xs

van Elswijk, G., Maij, F., Schoffelen, J.M., Overeem, S., Stegeman, D F., Fries, P. (2010). Corticospinal beta-band synchronization entails rhythmic gain modulation. J. Neurosci. 30(12), 4481–4488. https://doi.org/10.1523/JNEUROSCI.2794-09.2010

Wischnewski, M., Kowalski, G.M., Rink, F., Belagaje, S.R., Haut, M.W., Hobbs, G., Buetefisch, C.M. (2016). Demand on skillfulness modulates interhemispheric inhibition of motor cortices. J. Neurophysiol. 115(6), 2803–2813. https://doi.org/10.1152/jn.01076.2015

Witham, C.L., Wang, M., Baker, S.N. (2007). Cells in somatosensory areas show synchrony with beta oscillations in monkey motor cortex. Eur. J. Neurosci. 26(9), 2677–2686. https://doi.org/10.1111/j.1460-9568.2007.05890.x

Yanagisawa, T., Yamashita, O., Hirata, M., Kishima, H., Saitoh, Y., Goto, T., et al. (2012). Regulation of motor representations by phase-amplitude coupling in the sensorimotor cortex. J. Neurosci. 32(44), 15467–15475. https://doi.org/10.1523/JNEUROSCI.2929-12.2012

Zarkowski, P., Shin, C.J., Dang, T., Russo, J., Avery, D. (2006). EEG and the variance of motor evoked potential amplitude. Clin. EEG Neurosci. 37(3), 247–251. https://doi.org/10.1177/155005940603700316

Zrenner, C., Belardinelli, P., Müller-Dahlhaus, F., Ziemann, U. (2016). Closed-Loop Neuroscience and Non-Invasive Brain Stimulation: A Tale of Two Loops. Front. Cell. Neurosci.10, 92. https://doi.org/10.3389/fncel.2016.00092

Zrenner, C., Desideri, D., Belardinelli, P., Ziemann, U. (2018). Real-time EEG-defined excitability states determine efficacy of TMS-induced plasticity in human motor cortex. Brain Stimul. 11(2), 374–389. https://doi.org/10.1016/j.brs.2017.11.016

Zrenner, B., Zrenner, C., Gordon, P.C., Belardinelli, P., McDermott, E.J., Soekadar, S.R., et al. (2020). Brain oscillation-synchronized stimulation of the left dorsolateral prefrontal cortex in depression using real-time EEG-triggered TMS. Brain Stimul. 13(1), 197–205. https://doi.org/10.1016/j.brs.2019.10.007

